# Airway epithelial integrin β4 suppresses allergic inflammation by decreasing CCL17 production

**DOI:** 10.1101/513499

**Authors:** Lin Yuan, Xun Zhang, Ming Yang, Yizhou Zou, Yang Xiang, Xiangping Qu, Huijun Liu, Xizi Du, Leyuan Wang, Shuangyan Wu, Mengping Wu, Ling Qin, Qingwu Qin, Xiaoqun Qin, Chi Liu

**Affiliations:** Department of Physiology, Xiangya School of Medicine, Central South University, Changsha, Hunan, China; Centre for Asthma and Respiratory Disease, School of Biomedical Sciences and Pharmacy, Faculty of Health and Medicine, University of Newcastle and Hunter Medical Research Institute, Callaghan, New South Wales, Australia; School of Basic Medical sciences & Academy of Medical Science, Zhengzhou University, Zhengzhou, Henan, China; Department of Immunology, Xiangya School of Medicine, Central South University, Changsha, Hunan, China; Department of Respiratory Medicine, National Clinical Research Center for Respiratory Diseases, Xiangya Hospital, Central South University, Changsha, Hunan, China; Department of Internal Medicine, Division of Respiratory Disease, The Second Xiangya Hospital, Central South University, Changsha, Hunan, China

**Keywords:** ITGB4, airway epithelial cells, EGFR, CCL17, asthma, Th2 cells

## Abstract

Airway epithelial cells (AECs) play a key role in asthma susceptibility and severity. Integrin β4 (ITGB4) is a structural adhesion molecule that is downregulated in the airway epithelium of asthma patients. Specific ITGB4 deficiency in AECs induces exaggerated Th2 responses, severe allergen-induced airway inflammation and airway hyperresponsiveness (AHR) in mouse model of allergic asthma. However, the underlying mechanisms remain unexplored. In this study, we determine the role of ITGB4 of AECs in the regulation of Th2 response and in the induction of asthma and identify the underpinning molecular mechanisms. We found that ITGB4 deficiency led to exaggerated Th2 cells infiltration, inflammation and AHR and higher production of CCL17 in HDM treated mice. ITGB4-regulated CCL17 production in AECs was regulated by EGFR, ERK and NF-κB pathways. EFGR-antagonist treatment or the neutralization of CCL17 by antibody inhibited exaggerated pathological marks in HDM-challenged ITGB4-deficient mice. Together, these results demonstrated that ITGB4 of AECs negatively regulates the development of Th2 responses of allergic asthma by down-regulation of EGFR and CCL17 pathway.

## Introduction

Airway epithelial cells (AECs) form a barrier to environment hazardous stimuli by tight intercellular junctions and adhesion on basal membrane (Roche et al, 1993). Integrins play important roles in the adhesion, tissue repair and homeostasis of these cells (Pan et al, 2017). Among these molecules, integrin β4 (ITGB4) has been shown to regulate the adhesion of AECs on basal membrane through hemidesmosomal structure that is a specialized adhesion micro-structure attached to the extracellular matrix (Dowling et al, 1996). Previously, ITGB4 is implicated in the pathogenesis of allergic asthma, and its’ level is significantly decreased in AECs of asthma patients (Liu et al, 2010a; Xiang et al, 2014).

Allergic asthma is a chronic airway disorder that is characterized with epithelial desquamation, airway inflammation and airway hyperresponsiveness (AHR) to non-specific spasmogen (Matucci et al, 2018). Although CD4^+^ Th2 cells (Th2 cells) are widely recognized to orchestrate the development of the disease by expressing Th2 cytokines (e.g. interleukin (IL)-4, IL-5 and IL-13), there is compelling evidence that airway epithelium plays a vital role in the induction of aberrant immune responses underlying the pathogenesis of allergic asthma due to its’ frontline location to directly contact aerosol allergen and other environmental insultants (Lambrecht & Hammad, 2015). Indeed, AECs from asthma patients are often damaged and loss their homeostasis status including detachment, fragility and abnormal repair ability etc. (KleinJan, 2015; Wang et al, 2008). Upon allergen exposure, the repair process of epithelial layer of asthma patients is disrupted and is unable to restore the integrity of airway epithelial barrier (Georas & Rezaee, 2014; Heijink et al, 2014). As a result, lesion of airway epithelial barrier promotes the exposure of inhaled allergens to submucosal region whereby antigen presentation process is greatly accelerated. Furthermore, clinical studies have shown that high levels of epidermal growth factor receptor (EGFR) and transcription growth factor β (TGF-β) are produced by AECs of asthma patients (Boxall et al, 2006; Puddicombe et al, 2000), indicating the stressed repair process of airway epithelium. Mouse studies have also added to our understanding of how AECs can contribute to the pathogenesis of the disease. One example is that increased permeability of the airway epithelium to HDM exposure is associated with the heightened activity of nuclear factor of kappa B (NF-κB) and pro-inflammatory responses (Stacey et al, 1997). Indeed, a number of studies have shown that AECs direct subsequent immune responses through releasing various proinflammatory mediators to recruit and activate immune cells (Hallstrand et al, 2014; Holgate et al, 2009).

Infiltration of T lymphocytes and other immune cells is exquisitely modulated by pro-inflammatory mediators including chemokines (Castan et al, 2017). Distinctive patterns of chemokine receptors are differentially expressed on the surface of T helper cell subsets, among which several chemokine receptors drive the selective recruitment of Th2 cells into the asthmatic airways (Kim et al, 2001; Lukacs, 2001). For example, C-C chemokine receptor (CCR) 4 is preferentially expressed on Th2 cells, whose activation critically regulates the infiltration of these cells into the asthmatic airway upon allergen inhalation (Panina-Bordignon et al, 2001). Chemokine ligand (CCL) 3, 5, 17 and 22 are the main chemokines that selectively bind to CCR4 (Holgate, 2012). Indeed, high levels of CCL17 and CCL22 have been found not only in the lung and bronchoalveolar lavage fluid (BALF) of asthma patients but also in those of ovalbumin (OVA) sensitized and challenged mice (Gonzalo et al, 1999; Hijnen et al, 2004). CCL17 and CCL22 are predominantly produced by AECs and pulmonary macrophages, respectively (Gonzalo et al, 1999; Kawasaki et al, 2001). In particular, the former chemokine is increasingly being recognized as a key epithelial secreted attractant for Th2 cells recruitment to the asthmatic airways (Hijnen et al, 2004). Indeed, clinical studies have demonstrated that the level of CCL17 positively correlates with the degree of airway obstruction and fractional exhaled nitric oxide (FeNo) of asthma patients (Ying et al, 2005). However, the mechanisms how AECs contribute to the development of asthma by employing CCL17 still remain largely unknown.

Interestingly, ITGB4 is located at the basal surface of AECs and regulates the stable adhesion of these cells to the underlying basement membrane (Pan et al, 2016). ITGB4 has a unique long cytoplasmic domain subunit that could recruit a range of signaling molecules. ITGB4 plays a key role in the activation of several intracellular signaling pathways including phosphatidylinositol 3-kinase (PI3-K) (Shaw et al, 1997), extracellular signal-regulated kinases (ERK) 1/2 (Chen et al, 2010) and NF-κB (Nikolopoulos et al, 2004), indicating that this integrin potentially contributes to the instigation of subsequent immune response and inflammation. Although there is a correlation between ITGB4 and the induction of allergic asthma, much remains unknown about the underlying mechanisms. In particular, little is known about the contribution of ITGB4 pathway to Th2 regulated immune responses. Therefore, we sought to further unravel the role of such pathway in this study.

Using our well-established conditional knockout *in vivo* system, we were able to show that ITGB4 in AECs negatively regulates the development of Th2 responses and allergic asthma with a mouse model of house dust mite (HDM) induced asthma. We have also demonstrated a critical link between ITGB4 and CCL17 that underpins the activation of Th2 cells. We further explore this inflammatory paradigm through the administration of AG1478, a compound that has been demonstrated to bind antagonistically to EGFR (Wang et al, 2010), as a novel intervention strategy to disrupt the activation of ERK pathway and thus the Th2 responses. These findings highlight the critical role of ITGB4, EGFR and CCL17 in the regulation of Th2-inflammatory features and indicate a novel pathway that regulates key pathophysiological features of allergic asthma.

## Results

### Deficiency of ITGB4 in AECs aggravates the development of HDM induced allergic asthma

AECs specific ITGB4 conditionally knock out mice were treated with doxycycline to deplete this integrin as previously described (Fig 1A) (Liu et al, 2018). Mice were exposed HDM to induce allergic asthma. HDM exposure resulted in significantly increased airway reactivity and elevated levels of inflammatory cells in BALF and lung of ITGB4^+/+^ mice, as compared to PBS exposed control. Following HDM exposure, ITGB4^−/−^ mice exhibited significantly higher AHR than that of ITGB4^+/+^ mice (Fig 1B). HDM exposed ITGB4^−/−^ mice displayed significantly elevated histopathological scores, as compared to either PBS exposed ITGB4^−/−^ mice or HDM exposed ITGB4^+/+^ mice (Fig 1C). In line with aforementioned findings, the levels of BALF inflammatory infiltrates of HDM exposed ITGB4^−/−^ mice also augmented significantly when compared to those of HDM exposed ITGB4^+/+^ mice (Fig 1D). These observations reveal a negative role of ITGB4 in the regulation of the development of allergic asthma.

**Figure 1.**
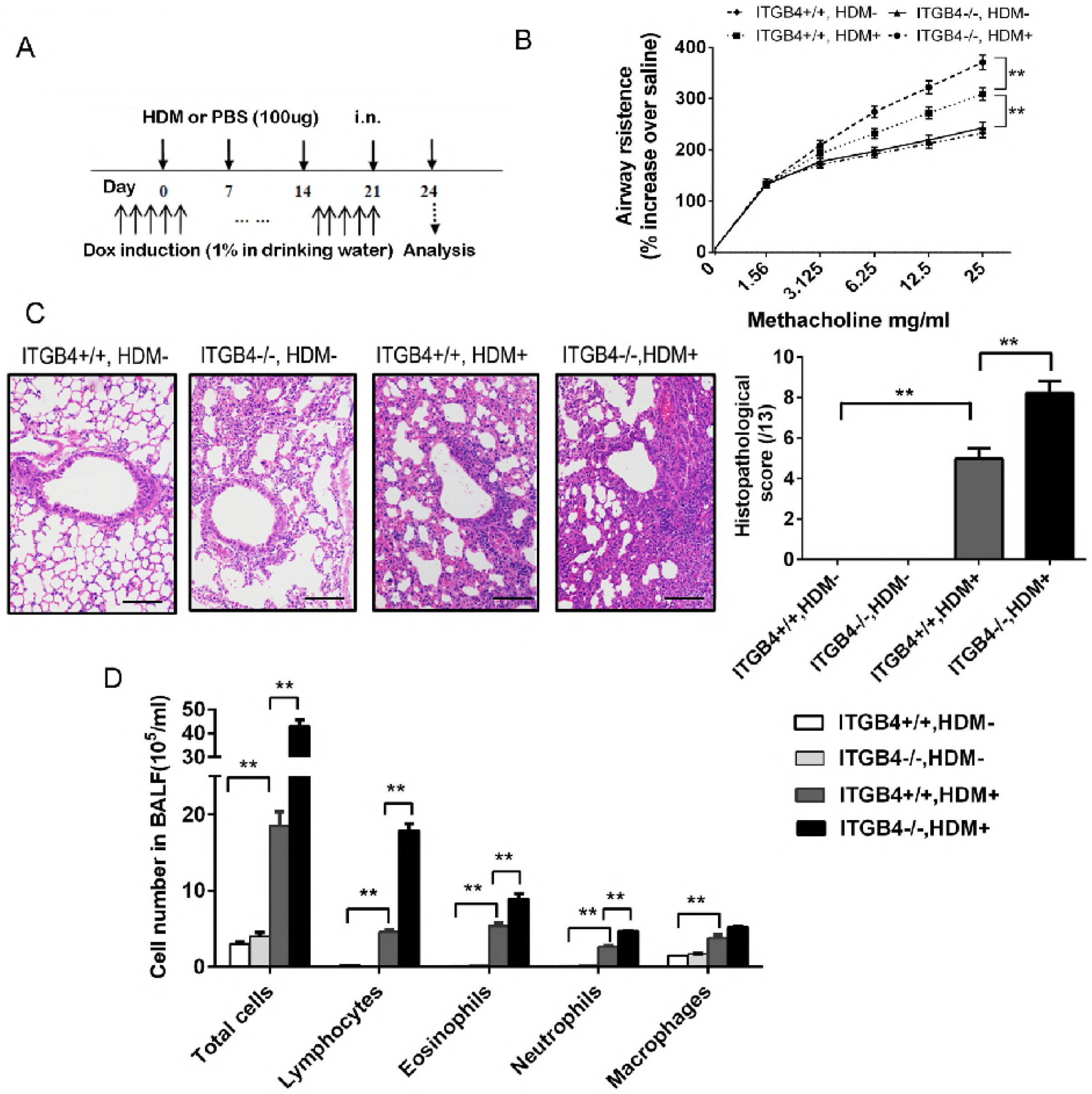
AHR and allergic disease of the lung is markedly exaggerated in the absence of airway epithelial ITGB4. A Mice were sensitized to HDM on days 0, 7, 14 or 21. Some mice were treated with 1% Dox in drinking water to specifically delete ITGB4 in airway epithelial cells. Control non-sensitized mice received PBS. B AHR was represented as airway resistance in response to methacholine. Data represent the mean ± SEM of six mice per group. **P < 0.01 by 2-way ANOVA followed by Fisher post hoc test. C Lung histology was assessed (n = 8), bars: 50 μm. Values represented as mean ± SEM. **P < 0.01 compared with controls using an unpaired, Student’s t test. D BALF inflammatory cell was counted (n = 10). Values represented as mean ± SEM. **P< 0.01 compared with controls using one-way ANOVA followed by Dunnett’s post hoc test.

### AECs-specific ITGB4 deficiency increases the numbers of Th2 and Th17 cells and the levels of their cytokines

To further characterize the impact of ITGB4 defect on the activation of T cells, we examined the infiltration of Th1, Th2 and Th17 cells by flow cytometry and the levels of their cytokines in lung by ELISA and qPCR, respectively. The levels of CD4^+^IL-4^+^ T cells, CD4^+^IL-13^+^ T cells and CD4^+^IL-17^+^ T cells increased significantly in the lung of both ITGB4^+/+^ and ITGB4^−/−^ groups following HDM exposure as compared to respective PBS control groups, and the levels of CD4^+^IFN-γ^+^ T cells did not display significant changes (Fig 2A). Of note, the infiltration of Th2 and Th17 cells were significantly greater in HDM exposed ITGB4^−/−^ animals than that of ITGB4^+/+^ animals. Likewise, the protein and transcript levels of IL-4, IL-5, IL-13 and IL-17A were significantly elevated as demonstrated in the BALF of HDM exposed ITGB4^−/−^ group, when compare to those of HDM exposed ITGB4^+/+^ group (Fig 2B and C). HDM exposure resulted in significantly increased levels of these cytokines in BALF and lung of ITGB4^+/+^ mice, as compared to those PBS exposed control. These data provide evidence that ITGB4 negatively regulates the activity of Th2 and Th17 cells but not Th1 cells.

**Figure 2.**
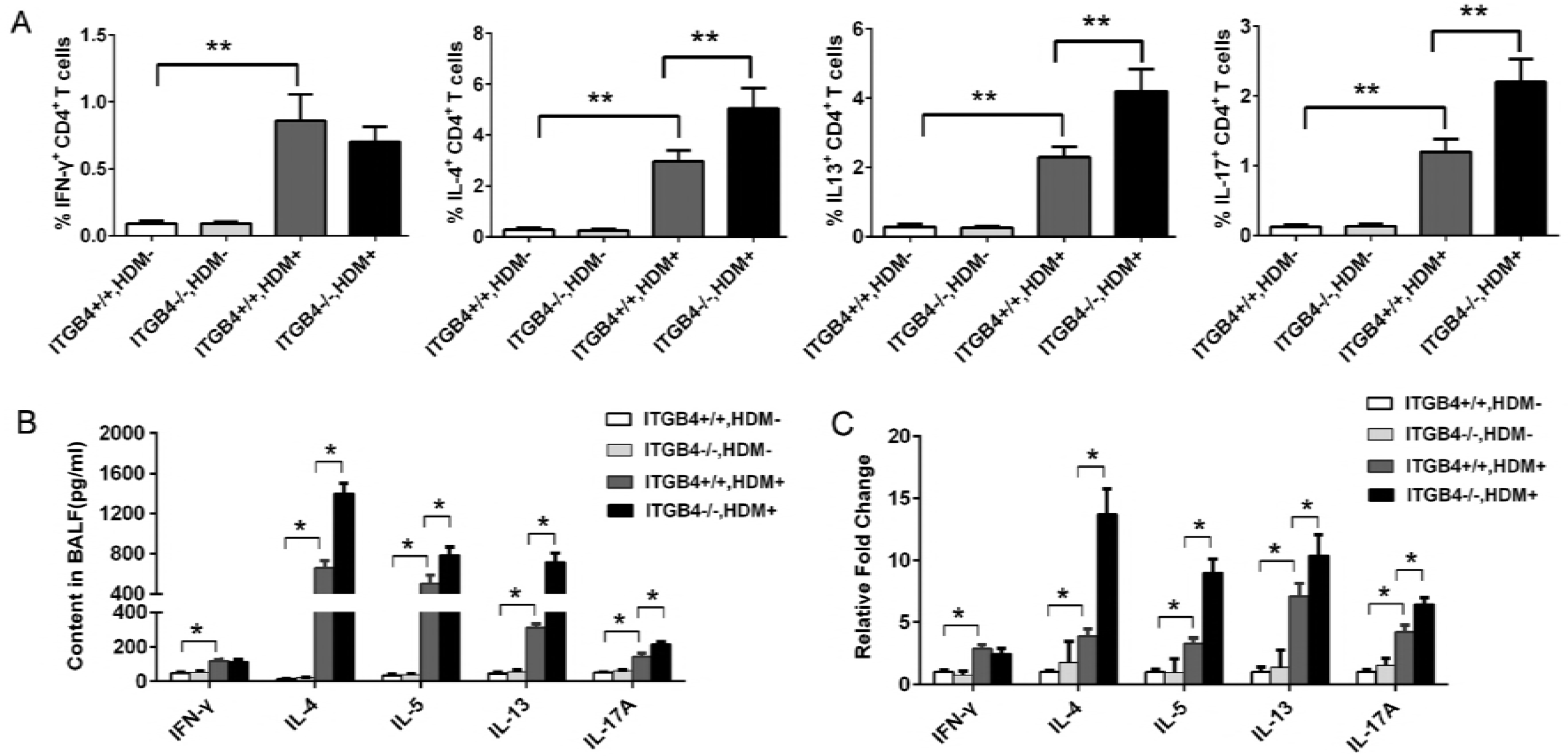
The infiltration T helper cells and the levels of BALF cytokine of ITGB4^+/+^ or ITGB4^−/−^ mice. A Mice were sensitized to HDM on days 0, 7, 14 or 21. Some mice were treated with 1% Dox in drinking water to specifically delete ITGB4 in airway epithelial cells. Control non-sensitized mice received PBS. Two days after the final challenge, the infiltration of IFN-γ^+^, IL-4^+^, IL-13^+^ and IL-17A^+^ T lymphocyte in lung of ITGB4^+/+^ and ITGB4^−/−^ mice was assessed by flow analysis (n = 12). Values represented as mean ± SEM. **P < 0.01 compared with controls using an unpaired, Student’s t test. B, C Two days after the final challenge, the levels of IFN-γ, IL-4, IL-13, and IL-17A protein in BALF (n = 8) and their transcripts in lung (n = 6) were examined by ELISA and qPCR, respectively. Values represented as mean ± SEM. *P< 0.05 compared with controls using one-way ANOVA followed by Dunnett’s post hoc test.

### ITGB4 negatively impacts on CCL17 production

Both clinic and animal studies have shown that Th2 cells orchestrate the development of allergic asthma (KleinJan, 2015). To determine how ITGB4 contributes the activation of Th2 cells, we examined the levels of CCL3, CCL5, CCL17 and CCL22 that could bind to CCR4, a Th2 chemotactic receptor (Assarsson et al, 2004; Qiu et al, 2018; Yoshie & Matsushima, 2015). Significant higher levels of CCL17 protein and transcript were found in the BALF and lung of HDM exposed ITGB4^−/−^ mice, as compared to those of HDM exposed ITGB4^+/+^ mice (Fig 3A and B). Interestingly, there was no significant difference in the transcript levels of CCL3, CCL5 and CCL22 in cultured AECs of HDM exposed ITGB4^−/−^ mice, as compared to those of HDM exposed ITGB4^+/+^ mice (Fig EV1). Sputum samples from patients with allergic asthma and healthy subjects were also collected to determine the secretion of CCL17. By using ELISA, levels of CCL17 were found significantly higher in the sputum of patients with allergic asthma, as compared to that of healthy controls (Fig 3C). Taken together, these findings indicate that ITGB4 in AECs critically regulates the expression of CCL17 and is associated with the pathogenesis of allergic asthma.

**Figure 3.**
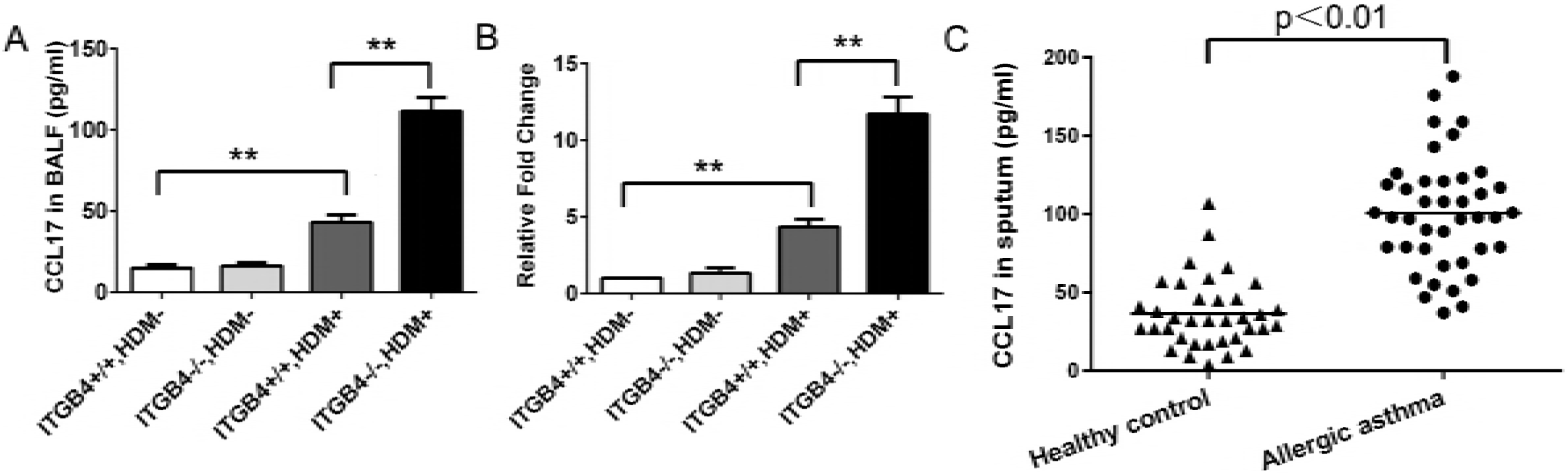
CCL17 production and Th2 cells infiltration are further elevated by the deficiency of ITGB4. A, B Mice were sensitized to HDM on days 0, 7, 14 or 21. Some mice were treated with 1% Dox in drinking water to specifically delete ITGB4 in airway epithelial cells. Control non-sensitized mice received PBS. CCL17 protein in BALF (n = 8) and the levels of CCL17 transcripts in lung (n = 6) were determined by ELISA and qPCR, respectively. Values represented as mean ± SEM. **P < 0.01 compared with controls using an unpaired, Student’s t test. C CCL17 expression detected in induced sputum samples from allergic asthmatic patients (n = 41) and control subjects (n = 35). Values represented as mean ± SEM. **P < 0.01 using an unpaired, Student’s t test.

### ITGB4 inhibits the EGFR phosphorylation of AECs

ITGB4 has been shown to regulate EGFR phosphorylation in liver cancer cells, breast cancer cells and gastric cancer cells, which is an important signalling mechanism for ERK activation (Bon et al, 2007; Huafeng et al, 2018; Leng et al, 2016). To determine whether ITGB4 regulates EGFR phosphorylation in our model, we treated isolated AECs from ITGB4^+/+^ and ITGB4^−/−^ mice with recombinant EGF to investigate the levels of EGFR phosphorylation. Basal level of EGFR phosphorylation in ITGB4^−/−^ AECs was higher than that in ITGB4^+/+^ AECs in the absence of EGF (Fig 4A). However, EGF significantly enhanced EGFR phosphorylation in ITGB4^−/−^ AECs, as compared to that in ITGB4^+/+^ AECs. Furthermore, assays with immunoprecipitation (Fig 4B) and immunofluorescence (Fig 4C) revealed a direct binding between ITGB4 and EGFR in cultured AECs. These results suggest that ITGB4 negatively regulates EGFR phosphorylation through physical association with the receptor.

**Figure 4.**
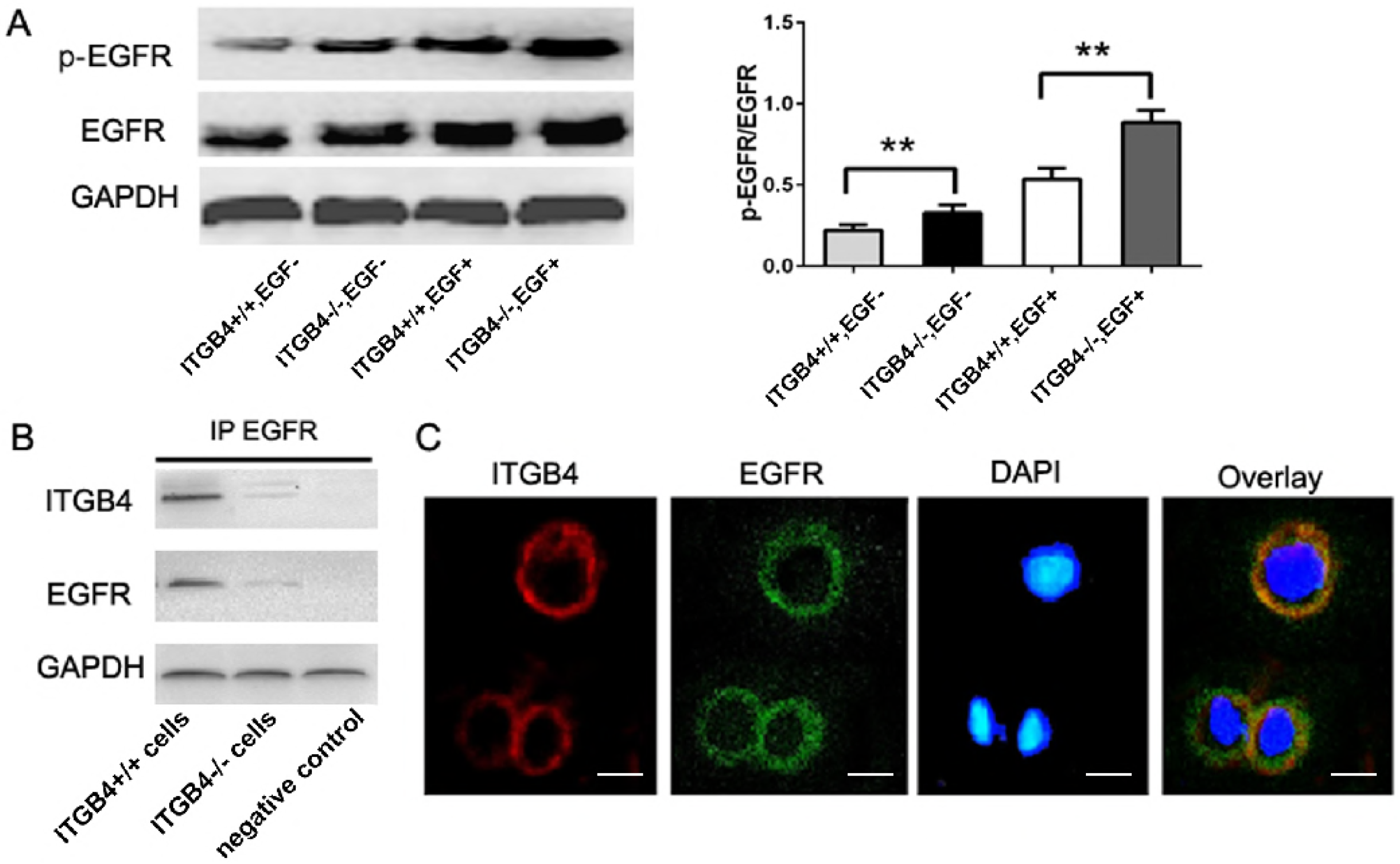
Physical interaction between ITGB4 and EGFR and the phosphorylation of EGFR in airway epithelial cells of ITGB4^+/+^ or ITGB4^−/−^ mice. A Airway epithelial cells were isolated from the lung of ITGB4^+/+^ or ITGB4^−/−^ mice. Cells were cultured and then stimulated with or without EGF (1 ng/ml) for 60 min. Western blot staining of stimulated AECs for p-EGFR and total EGFR, and p-EGFR/EGFR ratios normalized to vehicle control. Values represented as mean ± SEM for six samples from one experiment and representative of 3 independent experiments. **P<0.01 using an unpaired, Student’s t test. B Immunoprecipitation of stimulated AECs for interaction between ITGB4 and EGFR. C Immunofluorescence staining of stimulated AECs for co-localization of ITGB4 and EGFR, bars: 10 μm.

### ITGB4 regulates the production of CCL17 through the activation of EGFR, ERK and NF-κB pathways

To determine the role of EGFR mediated signaling pathways in ITGB4-regulated CCL17 production, we examined the expression of CCL17 in isolated AECs using multiple inhibitors, including AG1478 for EGFR phosphorylation, U0126 for ERK1/2 phosphorylation and SC75741 for NF-κB p65 activation. Western blot analysis revealed that treatments with AG1478, U0126 or SC75741 significantly inhibited the levels of p-EGFR, p-ERK1/2 or nucleus NF-κB p65 in ITGB4^−/−^ AECs than those in ITGB4^+/+^ AECs, respectively (Fig 5A, B and C). Importantly, these inhibitor treatments blocked heightened CCL17 production in ITGB4^−/−^ AECs. ERK activated pathways has been described to induce phosphorylation and translocation of the transcription factor NF-κB to the nucleus (37). Meanwhile, NF-κB inhibitors have been reported to block CCL17 expression in the alveolar epithelial cell line (Berin et al, 2001). To verify the possible involvement of NF-κB in the upregulation of CCL17, we studied p65 subunit of the active NF-κB complex upon stimulation with AG1478, U0126 or SC75741. As anticipated, treatments with AG1478, U0126 and SC75741 reduced the levels of nucleus NF-κB p65 whose binding sites are present in the CCL17 promoter in ITGB4^−/−^ AECs (Fig 5D). Together, these findings indicate that blocking EGFR, ERK and NF-κB pathways suppresses augmented CCL17 production in ITGB4^−/−^ AECs following HDM exposure.

**Figure 5.**
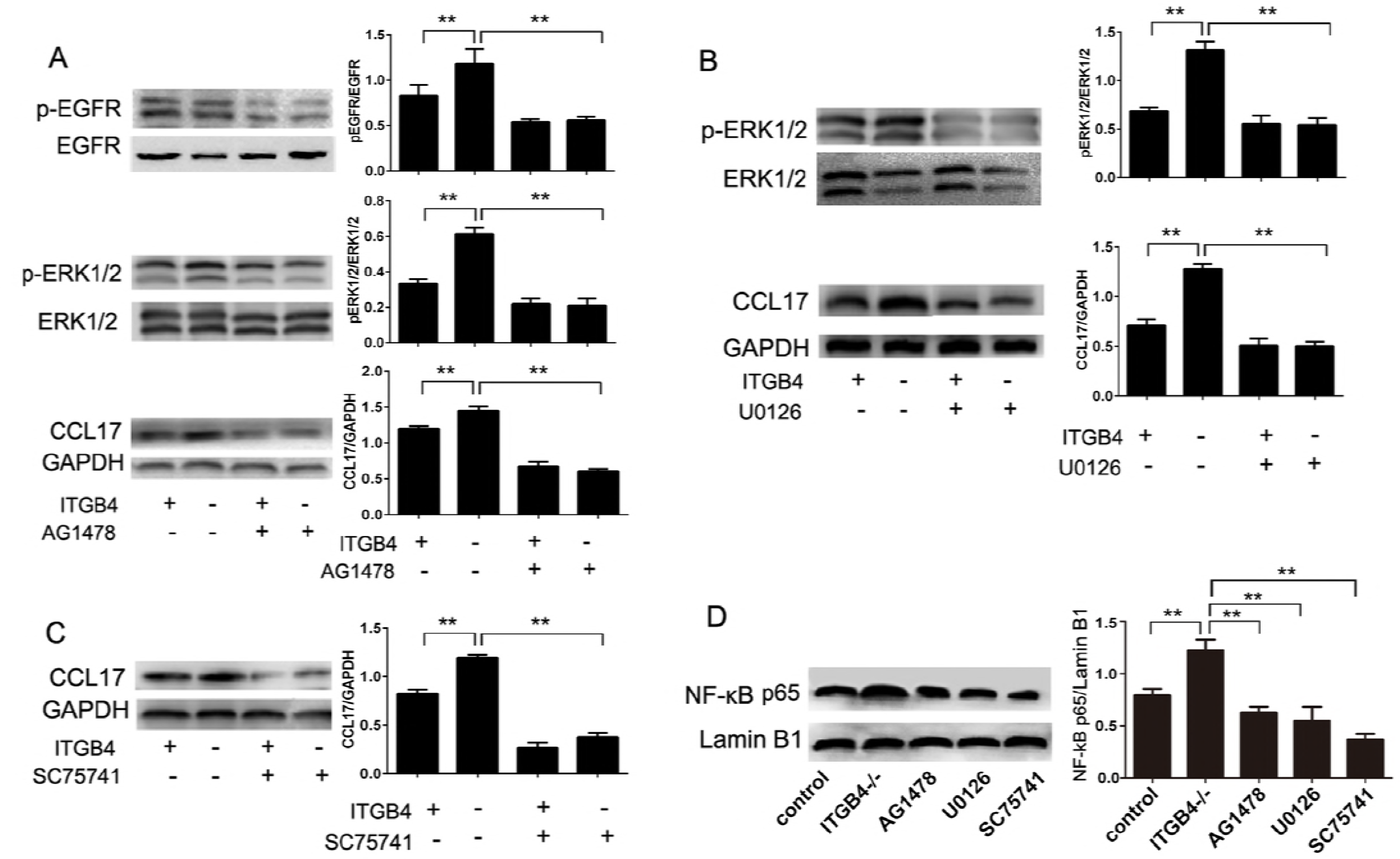
ITGB4 deficiency leads to augmented activation of EGFR, ERK1/2 and NF-κB pathways. A Airway epithelial cells were isolated from the lung of ITGB4^+/+^ or ITGB4^−/−^ mice. Cells were cultured and then stimulated with or without EGF (1 ng/ml) for 60 min. (A) Western blot staining of stimulated AECs for phosphorylated EGFR (p-EGFR), phosphorylate ERK1/2 (p-ERK1/2), total EGFR, total ERK1/2, and CCL17 in the presence or absence of EGFR tyrosine phosphorylation inhibitor AG1478. B Western blot staining of stimulated AECs for p-ERK1/2, total ERK1/2 and CCL17 in the presence or absence of ERK1/2 tyrosine phosphorylation inhibitor U0126. C Western blot staining of stimulated AECs for CCL17 in the presence or absence of NF-κB inhibitor SC75741. C Western blot staining of nuclear extracts of stimulated ITGB4^−/−^ AECs for NF-κB p65 in the presence or absence of AG1478, U1026 and SC75741, NF-κB p65/Lamin B1 ratios normalized to vehicle control. All values represented as mean ± SEM for six samples from one experiment and representative of 3 independent experiments. **P<0.01 using an unpaired, Student’s t test.

### Blockade of EGFR phosphorylation inhibits CCL17 production and exaggerated AHR, airway inflammation and Th2 cells infiltration

To investigate the role of EGFR signaling pathway in ITGB4-regulated CCL17 production, we blocked EGFR phosphorylation with AG1478 both *in vitro* and *in vivo* by using ITGB4^−/−^ AECs and mice. Administration of AG1478 inhibited the levels of p-EGFR in AECs, as compared to vehicle treatment (Fig 6A). Furthermore, inhibition of EGFR phosphorylation by AG1478 led to significantly decreased levels of CCL17 in cultured ITGB4^−/−^ AECs (Fig 6B). Exaggerated AHR and histopathological scores were also significantly reduced by EGFR blockade in HDM exposed ITGB4^−/−^ mice, as compared to those of vehicle treatment (Fig 6C and D). Flow cytometry revealed that AG1478 administration significantly reduced the levels of IL-4^+^ CD4^+^ T cells and IL-13^+^ CD4^+^ T cells but not IFN-γ^+^CD4^+^ T cells and IL-17^+^ CD4^+^ T cells in the lung of HDM treated ITGB4^−/−^ mice (Fig 6E). These data suggest that EGFR signaling pathway underpins ITGB4-regulated CCL17 production in AECs and thus contributes to the pathogenesis of Th2 inflammation.

**Figure 6.**
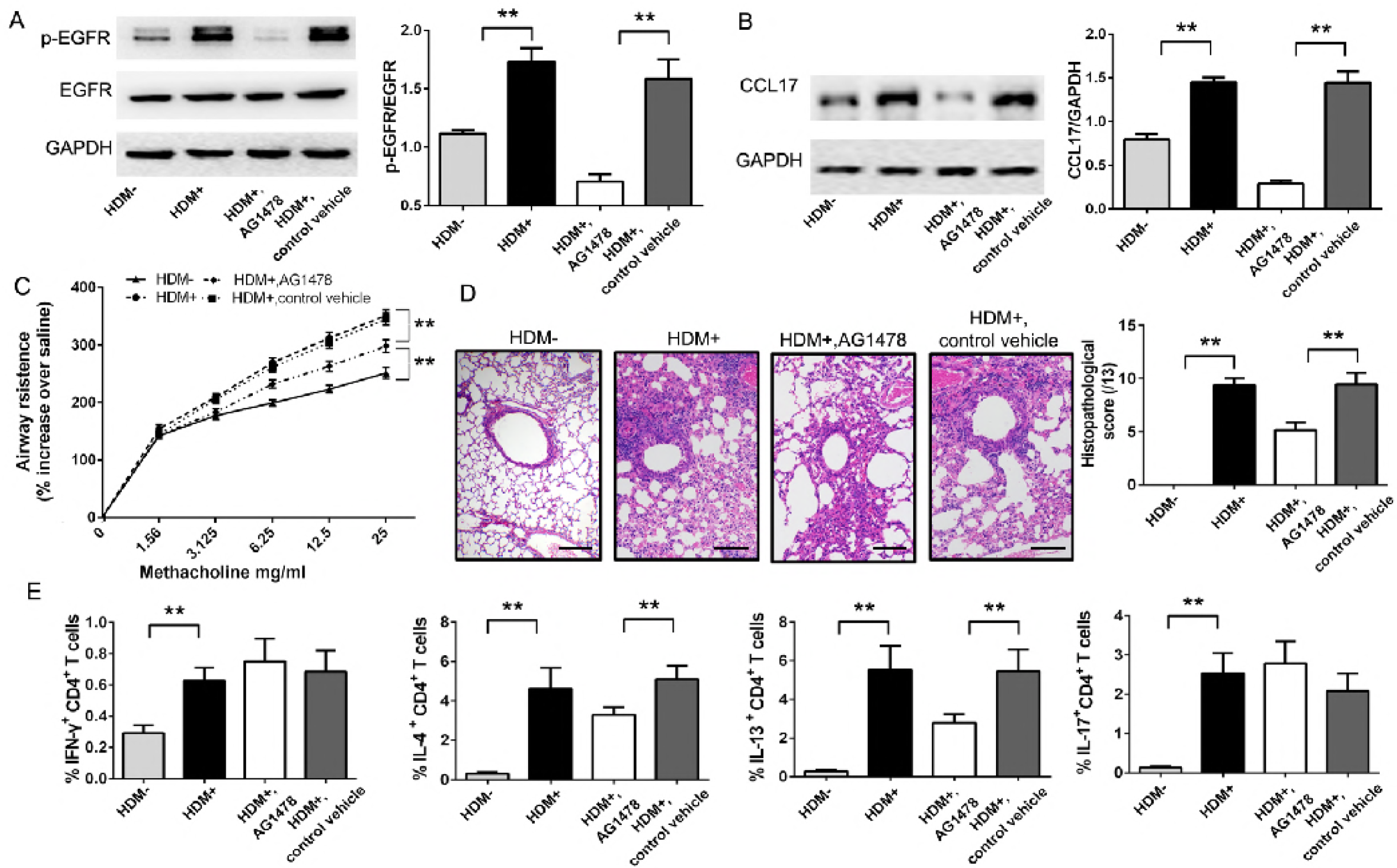
Blockade of EGFR phosphorylation inhibits both CCL17 production and subsequent enhanced AHR, lung inflammation in HDM exposed ITGB4^−/−^ mice. A, B AECs-specific ITGB4 conditional knock out mice were constructed and exposed HDM (HDM+) or PBS (HDM-) on days 0, 7, 14 or 21. AECs were isolated from these mice and some cells were treated with EGFR inhibitor AG1478 or control vehicle. The protein levels of EGFR, p-EGFR and CCL17 in AECs was detected by western blot. Values represented as mean ± SEM for 5 samples from one experiment and representative of 3 independent experiments. **P<0.01 using an unpaired, Student’s t test. Some mice received injection with either AG1478 or same volume of the control vehicle. C Two days after last HDM exposure, AHR was represented as airway resistance in response to methacholine. Data represent the mean ± SEM of 6 mice per group. **P < 0.01 by 2-way ANOVA followed by Fisher post hoc test. D, E Lung histology was assessed (n = 8) and the levels of IFN-γ^+^CD4^+^, IL-4^+^CD4^+^, IL-13^+^CD4^+^ and IL-17A^+^CD4^+^ T cells in lung were determined (n = 10), bars: 50 μm. Values represented as mean ± SEM. **P < 0.01 compared with controls using an unpaired, Student’s t test.

### Neutralization of CCL17 diminishes exaggerated AHR, airway inflammation and Th2 responses in HDM exposed ITGB4^−/−^ mice

As a key role for CCL17 is indicated in the pathogenesis, we neutralized this chemokine with anti-CCL17 mAb in HDM exposed ITGB4^−/−^ mice. Treatment with anti-CCL17 mAb significantly and greatly decreased the levels the chemokine in lung, as compared to isotype Ab treatment (Fig 7A). Neutralization of CCL17 significantly suppressed exaggerated AHR, airway inflammation in HDM exposed ITGB4^−/−^ mice (Fig 7B and C). Of note, anti-CCL17 mAb treatment reduced the levels of CD4^+^IL-4^+^ T cells and CD4^+^IL-13^+^ T cells but not CD4^+^IFN-γ^+^ T cells and CD4^+^IL-17^+^ T cells in the lung of HDM exposed ITGB4^−/−^ mice (Fig 7D). Similarly, neutralizing CCL17 decreased the protein and transcript levels of IL-4, IL-5, and IL-13, but not those of IFN-γ and IL-17A in the BALF and lung of HDM exposed ITGB4^−/−^ mice, as compared to isotype Ab treatment (Fig 7E and F).

**Figure 7.**
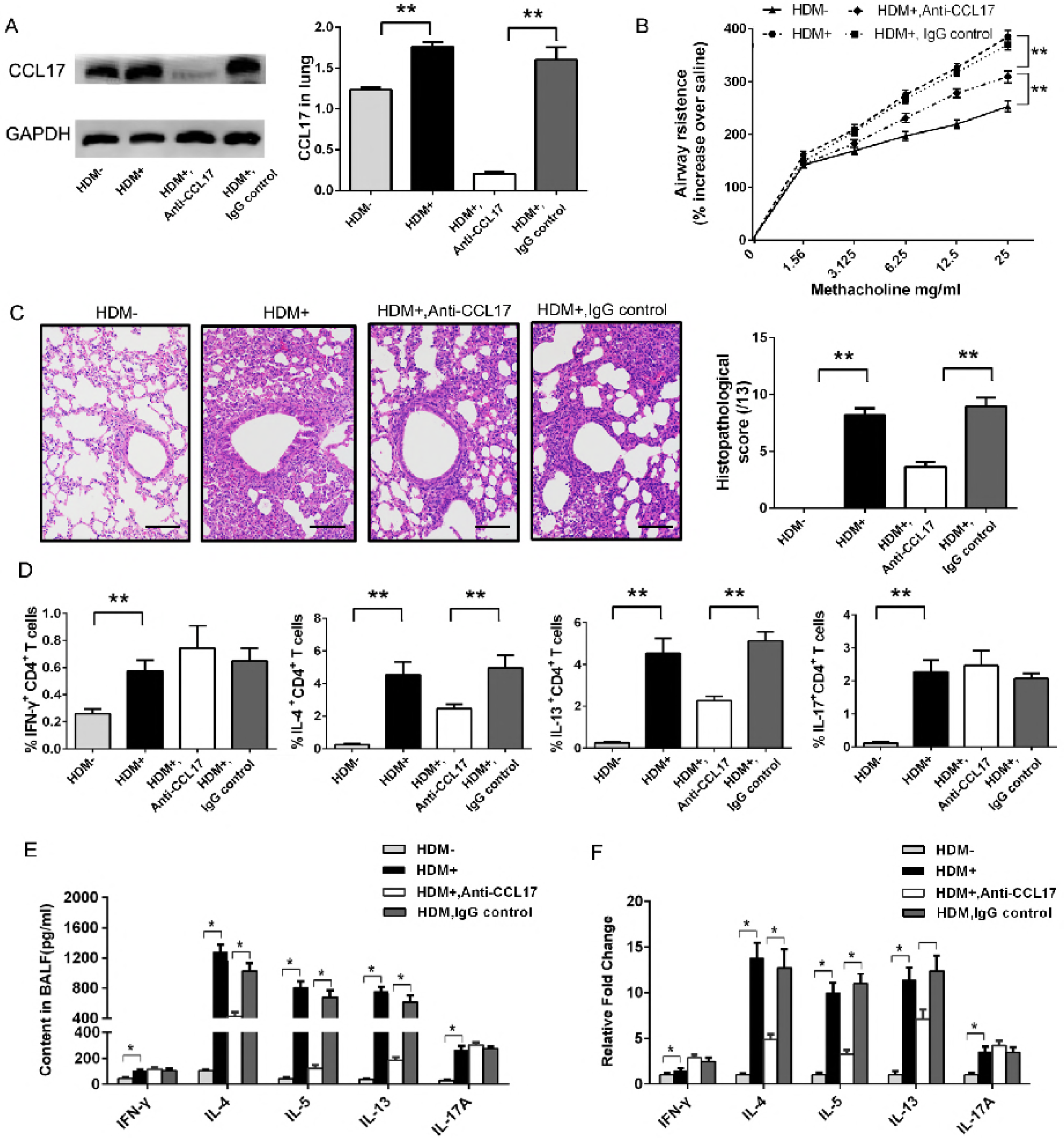
Neutralization of CCL17 reduces exaggerated AHR, airway inflammation and Th2 responses in HDM sensitized ITGB4^−/−^ mice. A Mice were treated with 1% Dox in drinking water to specifically delete ITGB4 in airway epithelial cells and were also sensitized with either HDM or PBS on days 0, 7, 14 or 21. Some mice received treatment with either anti-CCL17 or isotype control antibodies. Two days after the final challenge, CCL17 protein in lung was determined by western blot, ratios of CCL17/GAPDH normalized to vehicle control. Values represented as mean ± SEM for 6 samples from one experiment and representative of 3 independent experiments. **P<0.01 using an unpaired, Student’s t test. B Two days after last HDM exposure, AHR was represented as airway resistance in response to methacholine. Data represent the mean ± SEM of 6 mice per group. **P < 0.01 by 2-way ANOVA followed by Fisher post hoc test. C, D Lung histology was assessed (n = 8) and the levels of IFN-γ^+^CD4^+^, IL-4^+^CD4^+^, IL-13^+^CD4^+^ and IL-17A^+^CD4^+^ T cells in lung were determined (n = 10), bars: 50 μm. Values represented as mean ± SEM. **P < 0.01 compared with controls using an unpaired, 2-tailed Student’s t test. E, F Two days after the final challenge, the levels of IFN-γ, IL-4, IL-13, and IL-17A protein in BALF (n = 8) and their transcripts in lung (n = 6) were examined by ELISA and qPCR, respectively. Values represented as mean ± SEM. *P< 0.05 compared with controls using one-way ANOVA followed by Dunnett’s post hoc test.

## Discussion

Integrins are critical molecules for airway epithelial integrity and their expression is dramatically altered during chronic airway diseases such as allergic asthma (Huang et al, 1996; Qiu et al, 2017; Sundaram et al, 2017). However, the functional roles of these molecules in the regulation of aberrant immune responses and in the pathogenesis of disease are only beginning to be understood. We have previously reported that ITGB4 is downregulated in AECs of asthma patients and shown that it negatively regulates thymic stromal lymphopoietin (TSLP) production and antigen presenting process by DCs in an AECs-specific conditional knockout mouse system (Liu et al, 2018). In this study, we observed a negative regulatory role of ITGB4 in modulating the activity of Th2 cells through promoting CCL17 production in ITGB4^−/−^ mice, indicating an unidentified role of this integrin in the mechanisms underpinning the development of allergic asthma. Furthermore, our findings establish a new link between ITGB4, EGFR and CCL17 in AECs, and highlight the potential of targeting this pathway for the treatment of allergic asthma.

AECs are known as the first cell barrier to inhaled pollutants and allergen, and play a vital role in driving immune responses in respiratory disorders. Not only as the structural component that maintains epithelial architecture, integrins also play a key role in asthma pathogenesis. A recent study has shown that α5β1 integrin deficiency protects mice from cytokine-enhanced bronchoconstriction in a mouse model of asthma (Sundaram et al, 2017). However, it is difficult to specifically identify the mechanism for this protection, since integrin has been reported to regulate a wide range of biological processes by targeting intracellular pathways including FAK, Src, Ras, RhoA, EGFR, ERK, p53, etc. (Leng et al, 2016; Mitra & Schlaepfer, 2006; Stewart & O’Connor, 2015). In this context, our observation that the specific disruption of ITGB4 in AECs protects animal from allergic asthma makes this problem more tractable and suggests a central role of AECs in the induction of allergic asthma. In our study, exposure to HDM in the airways of ITGB4^−/−^ mice led to exaggerated airway inflammation and heightened AHR (Fig 1), which is in line with our previous findings (Liu et al, 2010b). We have also shown that the effects of ITGB4 deficiency in AECs are not just limited on pathophysiological changes in airway but also on the activation of T helper cells as AECs-specific ITGB4 negatively regulates the infiltration of Th2 and Th17 cells and the production of their cytokines (Fig 2). This observation suggests that ITGB4 contributes to the disease progress through the regulation of Th2 and Th17 activation.

The infiltration of T helper cells is exquisitely regulated by a wide range of chemokines (Castan et al, 2017; Guerreiro et al, 2011). While changes in levels of other CCR4-binding chemokines (e.g. CCL3, CCL5 and CCL22) did not show significant differences, the level of CCL17 was significantly elevated in the BALF and the lung of HDM exposed ITGB4^−/−^ mice (Fig 3). Notably, a significantly increased level of CCL17 was detected in the sputum of allergic asthma patients. Indeed, CCL17 plays a crucial role in the recruitment of Th2 cells into lung, as demonstrated by a mouse model of RSV infection in the lung (Monick et al, 2007). Several lines of evidence have shown that CCL17 can be produced by alveolar macrophages and dendritic cells as well (Ait Yahia et al, 2014; Staples et al, 2012). However, it is difficult to determine the exact roles of these immune cells in the regulation of CCL17. In this regard, our study is important in understanding the pathogenesis of allergic asthma, by demonstrating that the AEC-derived CCL17 is critical for the activation of Th2 cells.

Although the production of CCL17 by AECs is known, it is still largely obscure how integrins contribute to the increased expression of CCL17 in allergen induced asthma. With ITGB4^−/−^ mice, we were able to identify that the synchronization between ITGB4 and EGFR essentially orchestrates CCL17 production in AECs through the action of multiple downstream signaling pathways including EGFR, ERK1/2 and NF-κB (Fig 4-5). Studies in cancer research have shown that interaction between ITGB4 and EGFR phosphorylation has a significant impact on the progression and metastasis of hepatocellular carcinoma cells, mammary tumour cells and mammary tumour cells (Bon et al, 2007; Huafeng et al, 2018; Leng et al, 2016). EGFR, a receptor tyrosine kinase, can induce phosphorylation of multiple intracellular transcriptional regulators including ERK1/2, and PI3K pathway (Jiang et al, 2017; Zhu et al, 2015). Among these signaling regulators, EGFR is essential for the activation of ERK signaling that critically mediate the production of pro-inflammatory factors (Huang et al, 2017). Our results are in line with the aforementioned findings, and have identified that a direct interaction between ITGB4 and EGFR in airway epithelial cells drives the development of Th2 cells-associated allergic responses. Interestingly, AG1478 treatment completely blocked the phosphorylation of EGFR and reduced CCL17 production in the AECs of HDM exposed ITGB4^−/−^ mice to basal level. It also significantly inhibited the development of AHR, airway inflammation and the infiltration of Th2 cells (Fig 6). Furthermore, neutralizing CCL17 dramatically suppressed the manifestation of the disease in our model. Therefore, our data have shown that EGFR and CCL17 critically contribute to the disease, as demonstrated by its’ impact on Th2 cells infiltration, exaggerated AHR, airway inflammation in HDM exposed ITGB4^−/−^ mice.

Interestingly, we have also found that ITGB4 defect leads to heightened infiltration of Th17 cells and higher production of IL-17A (Fig 2). The importance of Th17 cells and their cytokines is implied not only in autoimmune diseases but also in allergic asthma (Chakir et al, 2003; Molet et al, 2001). Although EGFR antagonist or CCL17 mAb treatments significantly suppressed the migration of Th2 cells in HDM exposed ITGB4^−/−^ mice, they had no impact on Th17 cells infiltration. Furthermore, CCL17 mAb treatment did not affect IL-17A production. This may be related the increased TSLP production in AECs (Liu et al, 2018). Indeed, unaffected Th17 function is likely the reason why there exists residual AHR and airway inflammation after EGFR antagonist or anti-CCL17 mAb treatments in HDM exposed ITGB4^−/−^ mice. These observations indicate the differential and profound roles of ITGB4 in the induction of allergic asthma and highlight the complex and heterogeneous nature of allergic asthma. Furthermore, the infiltration of Th1 cells and the production of IFN-γ were not affected after EGFR-inhibitor or anti-CCL17 mAb treatments in HDM exposed ITGB4^−/−^ mice, suggesting a specific role of ITGB4 in the pathophysiology of allergic asthma by the regulation of Th2 and Th17 responses.

In conclusion, our data provide an insight that ITGB4 defect in AECs leads to elevated Th2 responses and exaggerated AHR and airway inflammation. By focusing on the role of ITGB4, we have developed models that allow dissection of the mechanisms predisposing to the development of Th2 responses, allergic inflammation and AHR. Here we demonstrate the importance of integrated signaling events between ITGB4, EGFR, ERK1/2 and NF-κB pathways specifically in AECs for the induction of allergic disease which may be clinically relevant. Understanding the contribution of this molecular network within AECs to the pathogenesis of allergic asthma may provide new therapeutic approaches for the treatment of the disease.

## Methods

### Collection of sputum samples

Asthmatic patients (defined by clinical diagnosis with evidence of bronchodilator reversibility testing, clinical assessment and a confirmed history of atopy with raised, allergen-specific IgE or positive radioallergosorbent test to at least one aero-allergen) (Matucci et al, 2018) were recruited and categorized by induced sputum inflammatory cell counts as allergic asthma. Participants provided written informed consent, approved by the No.20180308 Central south University Research Ethics Committees.

The demographic and clinical characteristics are shown in Table EV1. Induced sputum samples were obtained using nebulized hypertonic (4.5%) saline, treated with dithiothreitol and counts and viability (trypan blue exclusion) were determined by hemocytometer. The plugs were treated with a volume of dithiothreitol representing 4 times the mucus weight. Then, an equal volume of PBS was added to all samples before filtration with a 0.45 μm nylon filter. The supernatants were collected for CCL17 quantitation.

### Animals

Control wild type (WT) and AECs-specific ITGB4 conditionally knocked out (ITGB4^−/−^) mice were housed under barrier conditions in air-filtered, temperature-controlled units under a 12-hour light-dark cycle and with free access to food and water. The generation of AECs-specific ITGB4 conditionally knocked out mice was described previously (Liu et al, 2018). Briefly, to produce ITGB4^−/−^ mice, doxycycline (Dox; 1% in drinking water) was administered to 8-week-old CCSP–rtTA ^tg/-^/TetO-Cre ^tg/-^/ITGB4 ^fl/fl^ mice. All animal studies were approved by the No.201803079 Central South University at XiangYa Animal Care and Use Committee. All the methods were carried out in accordance with the relevant guidelines and regulations.

### Induction of allergic asthma and administration of anti-CCL17 neutralizing antibody and EGFR antagonist

For the induction of allergic asthma, mice were intranasally (i.n.) exposed to 100 μg HDM on days 0, 7, 14 and 21 as previous described (Draijer et al, 2013). Nonsensitized mice were i.n. treated with phosphate-buffered saline (PBS). On day 24, AHR was assessed, and then inflammatory infiltrates and histological changes in the lung were quantified as described previously (Nguyen et al, 2018). Some of ITGB4^−/−^ mice were intraperitoneally (i.p.) treated with 150 μg anti-mouse CCL17 monoclonal antibody (mAb, clone 110904, R&D Systems) or isotype control mAb (MAB006; R&D Systems, USA) in 200 μl PBS, twice a week for 3 weeks as previously described (Lee et al, 2018). Some of ITGB4^−/−^ mice were given intraperitoneal (i.p.) injections of EGFR inhibitor AG1478 (20mg/kg, Calbiochem, San Diego, CA) which was dissolved in CMC-Na (0.5%) or control vehicle three times weekly over the course of HDM treatment period (Wang et al, 2010).

### Measurement of lung function

Airway resistance was measured using a direct plethysmography (Biosystems XA; Buxco Electronics, USA), as previously described (Liu et al, 2010b). In brief, mice were anesthetized with a mixture containing xylazine (10 mg/kg) and ketamine (100 mg/kg) by i.p. injection (ratio 1:4, respectively; 150 μl per mouse). A cannula was then inserted into the trachea, and mice were ventilated with a tidal volume of 8 ml/kg at a rate of 145 breaths/min. Increasing doses of aerosolized methacholine (1.56, 3.125, 6.25, 12.5, 25mg/ml, Sigma-Aldrich) were delivered intratracheally (i.t.). Airway resistance was presented as percentage increase over baseline (saline challenge).

### BALF collection and cell counting

BALF was collected and processed as previously described (Nguyen et al, 2016). In brief, the lung was lavaged with 0.5 ml ice-cold PBS with 0.1 mM EDTA twice. Red blood cells were removed by using hypotonic red blood cell lysis buffer and BALF was then centrifuged to collect cellular infiltrate. Total cell numbers were quantified using a haemocytometer and cells were cytospun onto glass slides (ThermoFisher Scientific, Scoresby, Victoria, Australia). Differential leukocyte counts were determined based on morphological criteria by light microscopy (×100) on May-Grunwald and Giemsa-stained slides.

### Lung histology and immunohistochemical staining

Lungs lobes from mice were removed, fixed in 4% paraformaldehyde (5 ml), and embedded in paraffin blocks. Then, sections were stained with hematoxylin and eosin (H&E) (Sigma, St. Louis, MO) (Nguyen et al, 2018; Wang et al, 2013). Histopathological changes (inflammatory infiltrates) were scored blindly at morphological criteria, according to previous publications (Starkey et al, 2014).

Immunofluorescent staining was also performed on primary mouse AECs using the following antibodies: anti-ITGB4 (Ab182120, Abcam), anti-EGFR (ab52894, Abcam) in 1% BSA/PBS overnight at 4°C, washed with 1% BSA/PBS and incubated with Cy3-conjugated or FITC-conjugated goat anti-rabbit IgG (10 mg/ml; GE Healthcare) for 45 min respectively, then washed with 1% BSA/PBS and stained with 4’,6-diamidino-2-phenylindole (DAPI) (Sigma-Aldrich) for 10 min at RT. Slips were fixed on slides. ITGB4 and EGFR in AECs were visualized using a fluorescence microscope (Carl Zeiss MicroImaging GmbH, Göttingen, Germany) with a ×100 objective lens. Images were captured using a digital camera (Axio-Cam ICc3, Spectra Service, Ontario, NY) and analysed using AxioVision Rel. 4.7 software (Zeiss). Sampling was performed on eight to ten different areas for 40-60 cells of each slide.

### Flow cytometry

Single cell suspensions from lungs were prepared as previously described (Teijaro et al, 2011) and were used for flow cytometric analysis. In brief, cell suspensions were prepared by enzymatically digesting the lung tissue using dispase (Discovery Labware; Corning, Bedford, MA). Following incubation with surface marker antibodies anti-CD45, (clone 104, BD Biosciences, San Jose, CA), anti-CD3 (clone 17A2, BD Biosciences, San Jose, CA), anti-CD4 (clone GK1.5, BD Biosciences, San Jose, CA) and isotype controls (Rajavelu et al, 2015), cells were fixed using 4% PFA and permeabilized using Cytofix/Cytoperm solution (BD Biosciences, San Jose, CA). Cells were then incubated with anti-IFN-γ (Cat. no. 12-7311), anti-IL-4 (Cat. no. 12-7041), anti-IL-13 (Cat. no. 12-7133) and anti-IL-17 (Cat. no. 12-7177) antibodies (eBioscience, San Diego, CA). Numbers of positive cells were quantified by flow cytometry (FACSCanto flow cytometer, BD Biosciences, San Jose, CA). Data were collected on a FACSCanto flow cytometer and analysed with FlowJo software (version 7.6, Tree Star, Inc).

### ELISA assay

Levels of IFN-γ, IL-4, IL-5, IL-13, IL-13, IL-17 and CCL17 were determined with ELISA assays according to the manufacturer’s protocols (Sigma, St. Louis, MO, USA).

### RNA extraction, RT-PCR and quantitative RT-PCR

Total RNA was prepared using TRIzol reagent (Invitrogen) and phenolchloroform extraction from whole-lung tissues of mice and quantified on a SmartSpecTM Plus spectrophotometer (Bio-rad, USA) (Liu et al, 2012). cDNA was synthesized by RT-PCR using oligo d(T) primer (Invitrogen) on a T100 thermal cycler (Bio-Rad). Quantitative PCR (qPCR) was performed on a CFX96 Touch™ Deep Well Real-Time PCR Detection System (Bio-rad, USA) using TaqMan Gene Expression Master Mix (Applied Biosystems) with thermal cycling conditions. Primer sequences were described in Table EV2. Resulting mRNA levels were normalized to GAPDH and expressed as a fold-change relative to control samples.

### Western blot

AECs were isolated from mouse lung as previously described (Brockman-Schneider et al, 2008). CCSP (sc-365992, Santa Cruze) was used to sort CCSP^+^ AECs from ITGB4^−/−^ mice by flow cytometry. CCSP^+^ AECs were cultured and used as ITGB4^−/−^ AECs. Fifty μg of cell protein was separated by 10% SDS-PAGE and transferred to a polyvinylidene difluoride membrane. Some AECs were pretreated with AG1478 (1μM, an EGFR phosphorylation inhibitor), U0126 (10μM, an ERK phosphorylation inhibitor), or SC75741 (5μM, a NF-κB p65 inhibitor) respectively. Levels of ITGB4, CCL17, EGFR, phosphorylated EGFR (p-EGFR), ERK, phosphorylated ERK (p-ERK) and NF-B p65 were determined respectively with anti-mouse antibodies against these proteins by western blot as previously reported (Jiang et al, 2014). GAPDH and Lamin B1 were used as controls as indicated.

### Immunoprecipitation

Protein were extracted with radioimmunoprecipitation assay (RIPA) buffer: 50 mmol/L Tris-HCl, 150 mmol/L NaCl, 1% NP40, 0.5% sodium deoxycholate, 0.1%SDS, 5 mmol/L EDTA, pH=7.4. Protein concentration was measured by BCA protein determination kit (PC0020, Solarbio, China). Immunoprecipitations were performed overnight using the appropriate dilution of a polyclonal anti-ITGB4 antibody (Ab197772, Abcam) added to the lysates (1 μg proteins), under rotatry mixing for 1 at 4°C h. The immune complexes were allowed to bind to Protein A-Sepharose beads at 4°C for 3 h, washed (15 min, 4°C) with RIPA buffer and resuspended in loading buffer. Immunoprecipitates were subjected to SDS, 10% PAGE and immunoblotted on a polyvinylidene difluoride membrane. Immunocomplexes were stained with anti-ITGB4 (Ab197772, Abcam) as well as EGFR (Ab52894, Abcam) using ECL according to the manufacturer’s guidelines (NCI4106, Thermo Pierce, USA).

## Acknowledgements

We are gratefully to Professor Arnoud Sonnenberg for providing ITGB4^fl/fl^ mice and Professor Jeffrey A. Whitsett for providing CCSP–rtTA^tg/−^/TetO-Cre^tg/tg^ mice. This work was funded by grants #81570026, #81670002 and #3167188 from the NSFC; grants #2017JJ2402 from the Hunan Natural Science Foundation; grant #16K097, #14K109 from open Foundation of Hunan College Innovation Program; grants #2015QNRC001 from the Young Elite Scientists Sponsors hip Program by CAST and grant #2018zzts813, #2018zzts812 from the Fundamental Research Funds for the Central Universities of Central South University.

## Author contributions

CL, MY and XQQ conceived and designed this study. CL, LY, XZ, YX, XZD and YZZ carried out the experiments. YX, XPQ, HJL, LQ and QWQ contributed to the interpretation of the results. CL and MY wrote the paper. All authors provided critical feedback and helped shape the research, analysis and manuscript.

---

**Figure EV1.**
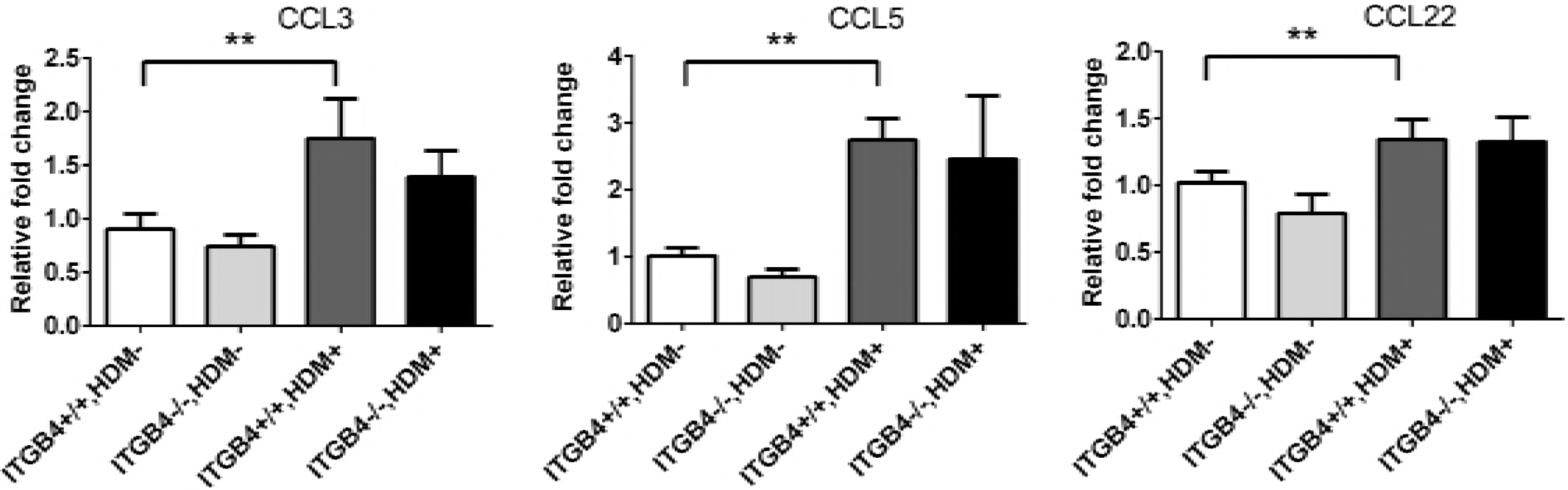
ITGB4 has no impact on the expression of CCL3, CCL5 and CCL22. ITGB4^+/+^ or ITGB4^−/−^ mice were exposed to HDM on days 0, 7, 14 or 21. The level of CCL22, CCL3 and CCL5 transcripts in airway epithelial cells (n = 10) were detected by qPCR. Values represented as mean ± SEM. **P< 0.01 compared with controls using controls using an unpaired, 2-tailed Student’s t test.

**Table EV1:** Characteristics of patients with allergic asthma and controls

|  | Control | Asthmatics |
| --- | --- | --- |
| Number | n = 35 | n = 41 |
| Gender (f/m) | 17/18 | 19/22 |
| Age | 27 (18-39) | 29 (16-45) |
| Atopy, n (%) ** | 7 (20) | 46 (71) |
| Total IgE (IU/ML)** | 65 (7-182) | 284 (29-1257) |
| FEV1(%)* | 105 (92–134) | 92 (45–114) |
| FEV1:FVC* | 91 (78–99) | 79 (51–86) |

FEV1, forced expiratory volume in 1s; FVC, forced vital capacity. Data are presented as mean±SD or median (range).

**Table EV2:**
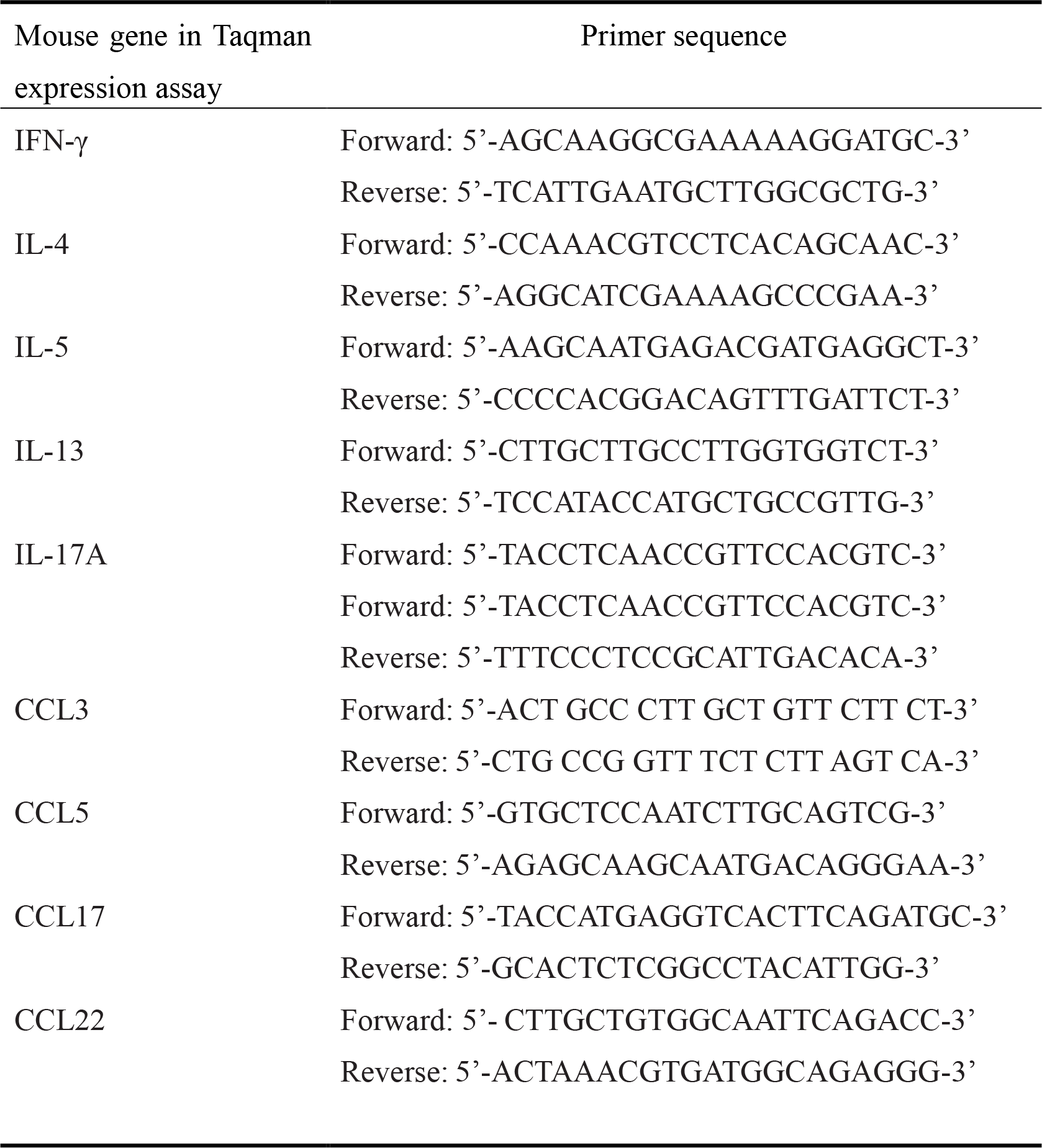
TaqMan primers for qRT-PCR.

